# Biotin-Independent *Saccharomyces cerevisiae* with Enhanced Growth: Engineering an Acetyl-CoA Carboxylase Bypass

**DOI:** 10.1101/2024.10.04.616611

**Authors:** Michaela Slanska, Sumbul A. Haider, Tsvetan Kardashliev, Dhanu Huck, Chang C. Liu, Thomas R. Ward

**Affiliations:** Department of Chemistry, University of Basel, and National Center of Competence in Research “Molecular Systems Engineering”, Basel, Switzerland; Department of Biomedical Engineering, University of California, Irvine, CA, USA

**Keywords:** *Saccharomyces cerevisiae*, biotin-independent, fatty acid synthesis, acetyl-CoA carboxylase, malonate, malonyl-CoA, metabolic engineering, biotechnology

## Abstract

Throughout evolution, most *Saccharomyces cerevisiae* strains have lost their ability to synthesize biotin, an essential cofactor of several carboxylating enzymes. As a result, the essential vitamin or its precursors must be uptaken from the environment and frequently supplemented in fermentations to achieve high cell densities. Engineering of a biotin-independent *S. cerevisiae* strain is of interest to eliminate the need for the external biotin supply. Herein, we describe the construction of a biotin-independent yeast strain by engineering a bypass of acetyl-CoA carboxylase, an essential biotin-dependent enzyme in the synthesis of fatty acids. Besides complete rescue of growth in biotin-free media, the resulting *S. cerevisiae* strains showed significantly improved growth on malonate compared to biotin. Beyond their industrial relevance, the yeast strains reported here can be valuable in areas of fundamental research, e.g., for developing a new selection marker or increasing the versatility of biotin-streptavidin technologies in living systems.

## 1. Introduction

*Saccharomyces cerevisiae* (brewer’s yeast) is one of the most commonly used microorganisms in biotechnological applications^1^. In addition to its use in the food and beverage industry, yeast is essential for producing bioethanol^2,3^ and potentially biodiesel^4^ as renewable energy sources. As such, the importance of *S. cerevisiae* will most likely continue to increase in the years to come. Up to 30% of the overall costs of fermented products arise from the materials used in industrial growth media. Biotin (vitamin B7) can be dfeicient in the media used for yeast fermentation (e.g., beet molasses) and thus needs to be supplied for unrestricted growth^5^. Therefore, engineering a biotin-independent *S. cerevisiae* strain is of interest to eliminate the need for an additional supply of this (expensive) vitamin.

Besides its industrial applications, *S. cerevisiae* is crucial in fundamental research as the simplest and most studied eukaryotic model organism^6^. Yeast has been extensively used to study eukaryotic cellular processes^7,8^ due to the ease of genetic manipulation through homologous recombination and the development of selectable auxotrophic marker genes^7,8^. In this context, identifying genes that could complement biotin auxotrophy would be interesting for developing a new selectable marker for genetic manipulations. Another potential of biotin-free yeast is in the biotin-streptavidin technology, which has been used in many research areas, e.g., as a detection system^9,10^, immobilization technique^11,12^, or to assemble artificial enzymes with new-to-nature reactivity^13,14^. However, applying the technology *in vivo* is complicated by endogenous biotin, which binds to the recombinantly-expressed streptavidin, competing with its binding to desired biotinylated molecules. Developing a biotin-free yeast strain would eliminate this interference, enhancing the utility of the biotin-streptavidin technology in living systems.

Biotin is a cofactor for several carboxylase-mediated reactions (**Figure 1a**)^15^. Acetyl-CoA carboxylase (ACC1 / HFA1) is the only essential enzyme in yeast that requires biotin as a cofactor. It catalyzes acetyl-CoA conversion to malonyl-CoA, an essential building block in fatty acid synthesis^16^. Pyruvate carboxylase isoforms PYC1 and PYC2 convert pyruvate to oxaloacetate^17^, which can replenish the TCA cycle or be used as an intermediate for gluconeogenesis. If grown in glucose-rich media where gluconeogenesis is inactive, this enzyme is not essential for yeast cells that rely on alcohol fermentation instead of the TCA cycle as their energy source. Biotin-dependent urea amidolyase DUR12 degrades urea to CO^2^ and NH^3^ but can be circumvented if other nitrogen sources are present^18^.

**Figure 1:**
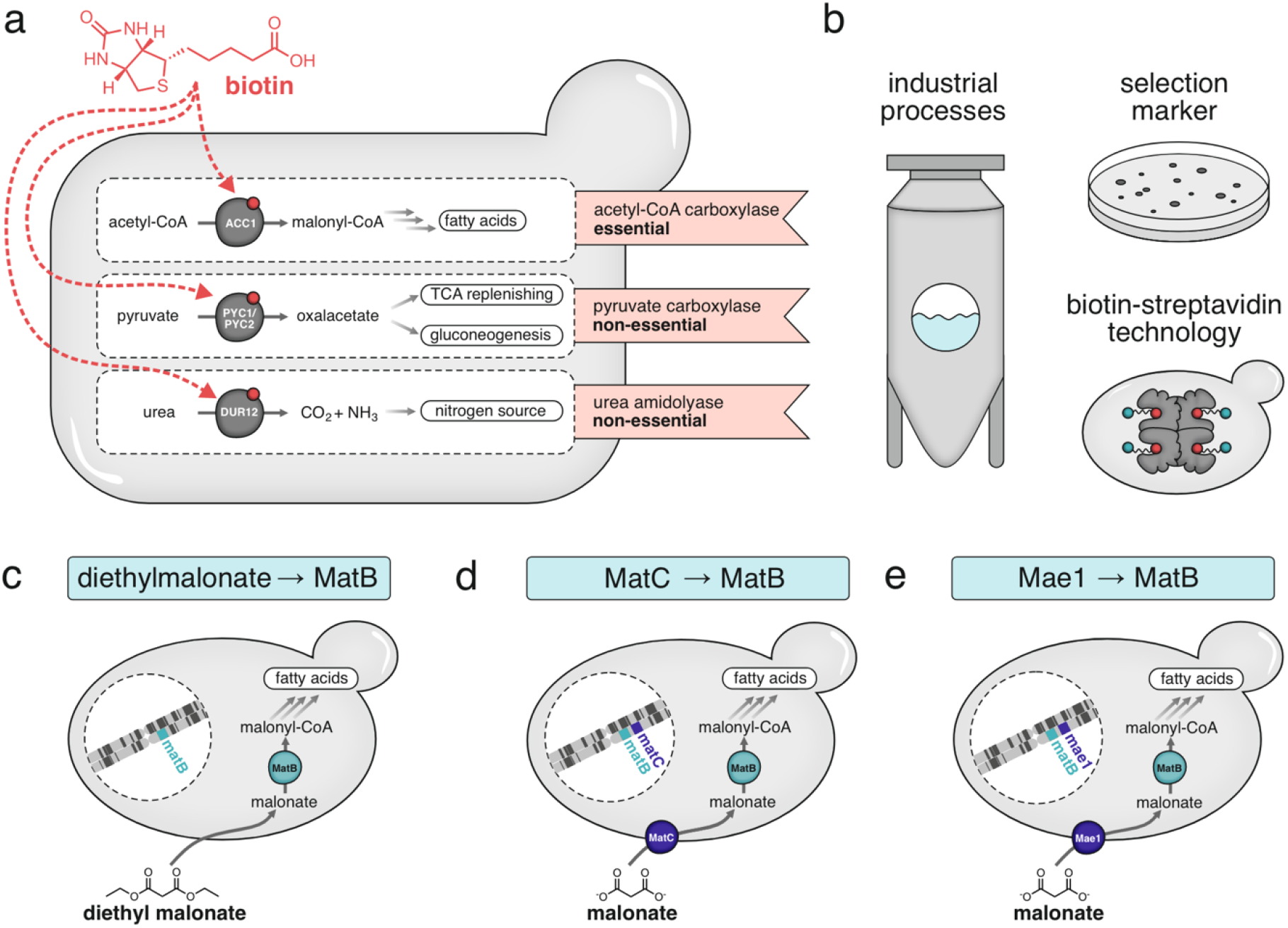
Malonyl-CoA synthesis bypass for biotin-independent *Saccharomyces cerevisiae*. **a)** Biotin-dependent enzymes of *S. cerevisiae* and their metabolic role. Acetyl-CoA carboxylase (ACC1) is the only essential enzyme that requires biotin as a cofactor. **b)** Potential applications of biotin-independent yeast in industry and fundamental research. **c)** An ACC1 bypass utilizing cell-permeable diethylmalonate and malonyl-CoA synthetase from *Rhizobium leguminosarium* bv. *Trifolii* (MatB). **d)** An ACC1 bypass utilizing the dicarboxylate carrier-protein from *Rhizobum trifolii* (MatC) and MatB. **e)** An ACC1 bypass utilizing the dicarboxylate plasma membrane transporter from *Schizosaccharomyces pombe* (Mae1) and MatB.

This work describes the engineering of a biotin-independent strain of *S. cerevisiae* to facilitate the abovementioned applications for large-scale industrial fermentations and fundamental research (**Figure 1b**). While efforts have previously been reported to engineer biotin-prototrophic strains of yeast^19,20^, this is, to the best of our knowledge, the irst instance where a fully biotin-independent strain has been constructed by bypassing ACC1-catalyzed malonyl-CoA synthesis. Surprisingly, the resulting engineered *S. cerevisiae* strains displayed not only complete rescue of growth but also improved growth on malonate compared to biotinylated media. Since the engineered malonate-dependent fatty acid synthesis pathway outperformed the endogenous ACC1 pathway, the work reported here may be of interest to enhance the productivity of industrial yeast applications and reduce the associated costs of yeast fermentations.

## 2. Results and discussion

### 2.1 Engineering of biotin-independent strains of *S. cerevisiae*

*S. cerevisiae* generally harbor only a partial biotin synthesis pathway (except for specific yeast strains, e.g., sake yeast), rendering their growth dependent on externally added biotin or biotic precursors, KAPA, DAPA, or DTB^21–23^. We hypothesized that the need for this vitamin may be eliminated by bypassing the only essential biotin-dependent yeast enzyme, ACC1, which catalyzes the conversion of malonate to malonyl-CoA^16^.

To create a biotin-independent yeast, an alternative pathway to malonyl-CoA was selected: malonyl-CoA synthetase, MatB from *Rhizobium trifolii*^24^, catalyzing the conversion of malonate to malonyl-CoA. This reaction could substitute ACC1 in fatty acid synthesis, but only in conjunction with an efficient malonate-uptake system.

Since malonate contains two negatively charged carboxylate groups, its transport across biological membranes is limited in the absence of a dedicated membrane channel. To circumvent this problem, we evaluated three strategies to facilitate its uptake. In the first approach, we supplemented diethylmalonate to the growth medium (**Figure 1c**). We hypothesized that the neutral compound may easily cross the cell membrane and would be subsequently hydrolyzed by endogenous carboxylesterases to yield malonate as substrate for MatB. As a second strategy, inspired by a previously-engineered biotin-independent *Escherichia coli* strain^25^, we introduced a dicarboxylate carrier protein MatC from *Rhizobium trifolii*^24^ to facilitate malonate uptake (**Figure 1d**). Lastly, we replaced this transporter with a dicarboxylate transporter Mae1 from the yeast *Schizosaccharomyces pombe*^26^, previously reported to facilitate malonate uptake into *S. cerevisiae*^27^ (**Figure 1e**).

We engineered a *S. cerevisiae* strain derived from strain F102-2 by encoding the appropriate genes (*matB, matC, mae1*) in its genome (**Figure 2a**). We used promoters of varying strengths^28^ to test the effect of different expression levels of each protein on cell growth. Eleven strains were constructed and genomic integrations were verified by colony-PCR (**Supplementary Figure 1**). Before the growth of the strains on (diethyl)malonate was tested, residual biotin was eliminated by serial passaging of the engineered strains in biotin-free media until minimal growth was observed (**Supplementary Figure 2**).

**Figure 2:**
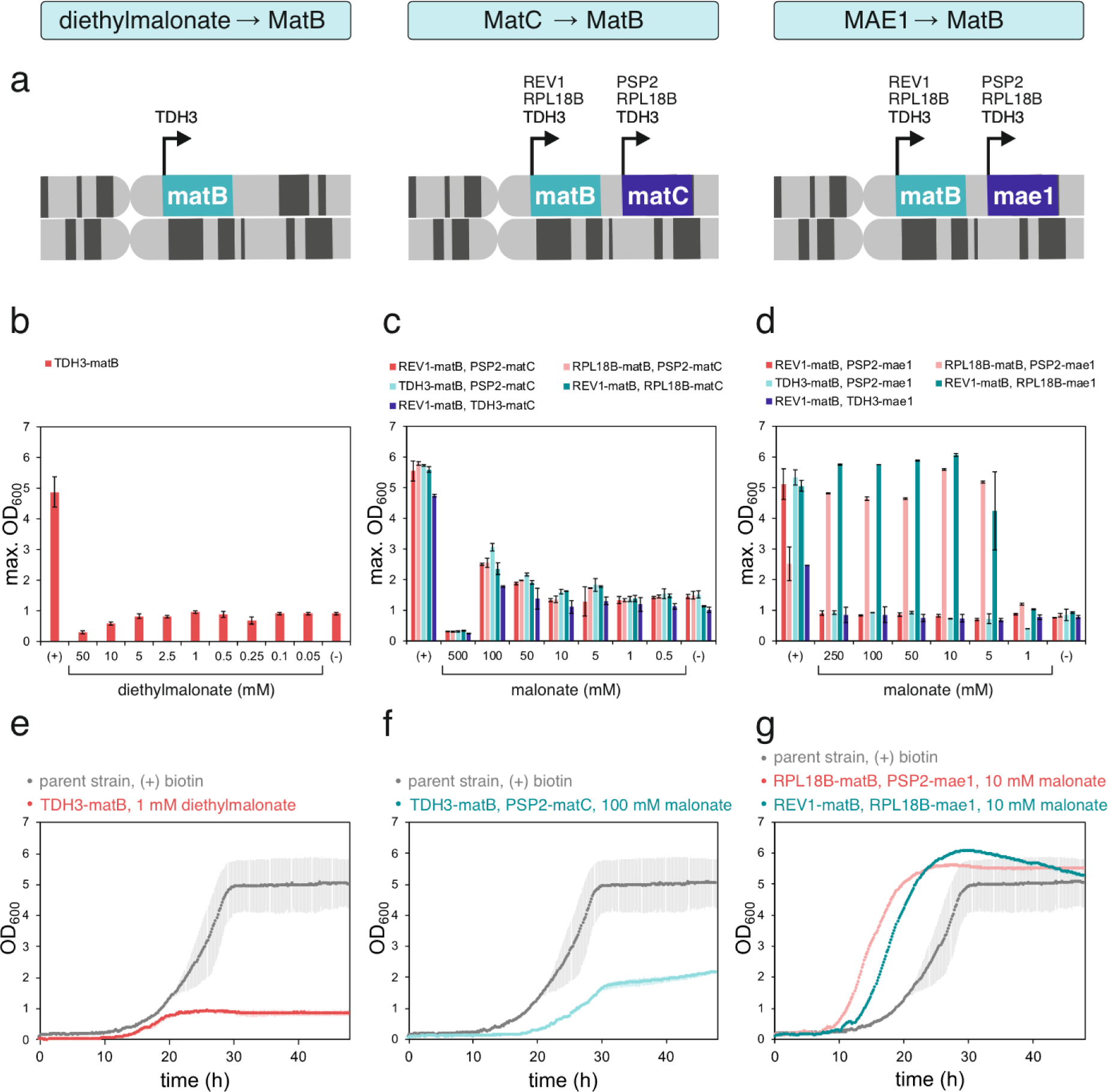
Engineering biotin-independent strains of *Saccharomyces cerevisiae*. **a)** Genes encoding MatB, MatC, and Mae1 proteins under the control of weak (REV1, PSP2), medium (RPL18B), or strong (TDH3) promoters were integrated into the Int. 1 MYT locus of Chromosome I. **b**-**d)** Growth-dependence of engineered strains on (diethyl)malonate concentration series. (+) – 2 *μ*g L^-1^ biotin, (-) – without biotin and (diethyl)malonate. **e**-**g)** The growth of the best-performing strains at their optimal (diethyl)malonate concentrations for each bypass strategy compared to the growth of the parent strain (without *matB, matC*, or *mae1*) on biotin (2 *μ*g L^-1^, positive control).

### 2.2 ACC1 bypass utilizing diethylmalonate and MatB

To test the import strategy utilizing uncharged diethylmalonate, we initially engineered a strain with high *matB* expression under the control of a strong TDH3 promoter. We tested its growth-dependence on diethylmalonate concentration series and compared it to growth on biotinylated media as a positive control (**Figure 2b, Supplementary Figure 3**). However, we did not observe significant growth complementation at micromolar concentrations of diethylmalonate. The engineered strain performed similarly to the parent strain, exhibiting only limited growth in the absence of biotin. At concentrations of diethylmalonate in the millimolar range, a slight improvement in growth was observed, with the best growth detected at 1 mM diethylmalonate (**Figure 2e**). However, even under these conditions, the growth-enhancement was minimal (19% cell density reached compared to media supplemented with biotin). Diethylmalonate concentrations exceeding 2.5 mM hindered growth. This could be explained by using ethanol, which can be toxic at higher contents, as a cosolvent for the water-insoluble diethylmalonate, preventing us from fully optimizing the supply. Accordingly, we did not test strains with different expression levels of *matB*. Instead, we shifted focus to the strategies relying on malonate, as it is water-soluble and does not require the use of toxic cosolvents.

### 2.3 ACC1 bypass utilizing MatC transporter and MatB

At neutral pH, malonate is charged, and thus soluble in water, eliminating the need for cell-toxic co-solvents. However, due to this property, it cannot permeate the yeast’s cell membrane, thus requiring a specific transporter for its uptake. We took inspiration from *Jeschek et al*., who previously engineered a biotin-independent strain of *Escherichia coli*^25^ by expressing the MatC transporter from *R. trifolii* on its membrane. We integrated the *matC* gene into the genome of *S. cerevisiae*, constructing five strains with varying MatB and MatC expression levels, and tested their growth on malonate concentration series (**Figure 2c, Supplementary Figure 4**).

A clear dependence of growth on malonate concentration was observed in all engineered strains (expressing MatB and MatC), which was not observed in the parent strain. This confirms the feasibility of substituting the biotin-dependent ACC1 by a malonate bypass. The best growth was observed in the strain with high MatB expression (TDH3 promoter) and low MatC expression (PSP2 promoter) (**Figure 2f**) in the presence of 100 mM malonate. However, even at such a high concentration, full growth complementation could not be achieved (compared to the parent strain growth on biotin). At higher concentrations than 100 mM, the increased osmotic pressure caused cell toxicity.

Furthermore, increasing the expression of the MatC transporter hindered the strains’ growth in biotin-containing media (**Supplementary Figure 5**), highlighting its toxicity to yeast. This was not surprising, as heterologous expression of foreign membrane proteins often leads to toxicity^29^. We thus set out to replace MatC for an alternative malonate transporter from the yeast *Schizosaccharomyces pombe*, Mae1.

### 2.4. ACC1 bypass utilizing Mae1 transporter and MatB

We constructed five strains with varying expression levels of MatB and Mae1 and evaluated their growth-dependence on malonate concentration (**Figure 2d, Supplementary Figure 6**). Three of the five strains displayed no growth improvement with malonate supply. The remaining two strains (RPL18B-*matB*, PSP2-*matC*, and REV1-*matB*, RPL18B-*matC*) were not only able to rescue growth when grown on malonate but, strikingly, surpassed the growth in biotinylated media (**Figure 2g**). Both strains (**Figure 3a**) proved biotin-independent at ∼5 mM malonate, a much lower concentration than required for the MatC transporter. The maximum OD^600^ and the maximum growth rate were significantly improved over the parent strain (**Figure 3b**), highlighting the role of MatB and Mae1 proteins in substituting the role of ACC1.

**Figure 3:**
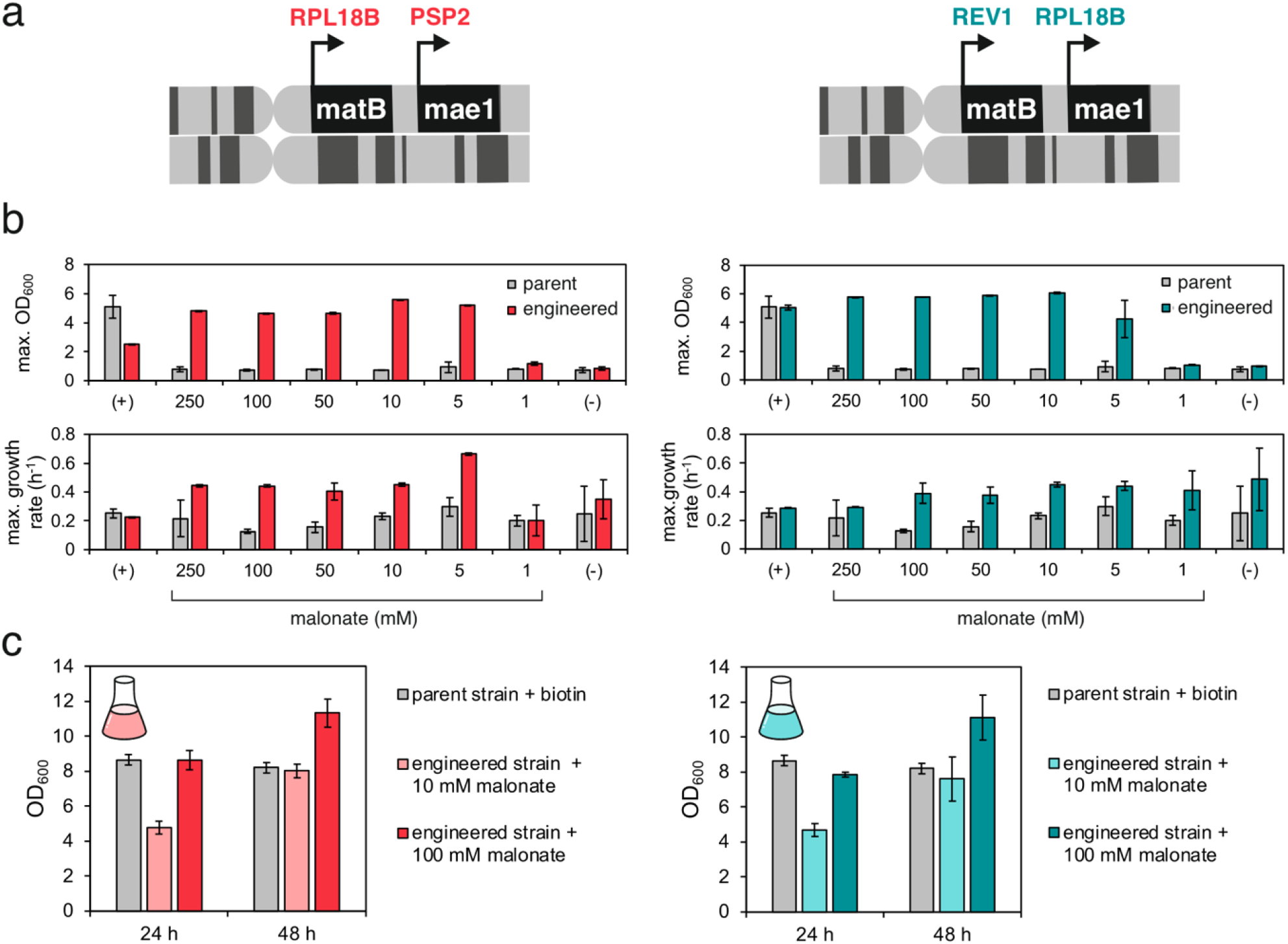
Two engineered strains containing Mae1 and MatB display complete rescue of growth on malonate. **a)** Two strains with complete rescue of growth on malonate: RPL18B-*matB*, PSP2-*mae1*, and REV1-*matB*, RPL18B-*mae1*. **b)** The engineered strains display malonate-dependent growth in biotin-free media. (+) – 2 *μ*g L^-1^ biotin, (-) – without biotin and malonate. **c)** Shake-lask experiments conirm that the engineered strains reach growth complementation and further improve growth over the parent strain on biotin (2 *μ*g L^-1^).

To validate the results obtained in the 96-well plate format used for growth-curve measurements, a shake-lask cultivation was carried out, where the growth of the two biotin-independent strains on 10 mM and 100 mM malonate was compared to the growth of the parent strain on biotin (**Figure 3c**). Once again, after 48 hours, both strains reached higher cell density when grown on 100 mM malonate compared to biotinylated media. Unlike certain bacteria^30,31^, yeast cannot utilize malonate as an energy source through decarboxylation to acetate. Therefore, this improvement in growth highlights that the engineered malonate-dependent pathway likely outperforms the endogenous ACC1 pathway in fatty acid synthesis. Such faster-growing *S. cerevisiae* could be of interest for enhancing productivity in industrial yeast applications, such as biofuel or biopharmaceutical production.

## 3. Conclusions

Most *Saccharomyces* cerevisiae strains, including common laboratory strains, have lost their ability to synthesize biotin through the course of evolution^21,32^. Consequently, as an essential cofactor for fatty acid biosynthesis, this vitamin must often be supplied in the media used for industrial fermentations^5^. In this study, we engineered two biotin-independent yeast strains with an alternative fatty acid synthesis pathway by bypassing ACC1-catalyzed malonyl-CoA synthesis. The strains express a dicarboxylate plasma membrane transporter from *Schizzosaccharomyces pombe* Mae1 and a malonyl-CoA synthetase from *Rhizobium leguminosarium* bv. *Trifolii* MatB. They grow in biotin-free media in the presence of malonate, a cheap chemical abundant in many plant species^33–35^.

In addition to complete rescue of growth, the engineered yeast reached higher cell densities when grown on malonate compared to biotin. Accordingly, the engineered malonate-dependent pathway liberates the cells from their biotin dependency and, most importantly, outperforms the natural malonyl-CoA synthesis route. Therefore, the fatty acid synthesis bypass presented herein may be of interest for enhancing productivity in industrial applications of *S. cerevisiae*.

Besides their applications in industry, the strains presented here hold great value in several areas of fundamental research. As an example, allowing yeast to grow in a biotin-free environment allows the biotin-streptavidin technology to be applied *in vivo* in the cytosol of *S. cerevisiae* without the interference of endogenous biotin. This technology is very promising, for instance, in the area of artificial metalloenzymes, where biotinylated metal complexes can be anchored to streptavidin to create enzymes with new-to-nature reactivity^36,37^. Incorporating streptavidin-based artificial metalloenzymes into *S. cerevisiae* could introduce novel chemistries into its metabolic pathways, potentially enabling more sustainable biotechnological production of chemicals that are otherwise unattainable through natural cellular processes^37^.

## 4. Materials and Methods

### 4.1 Suppliers

Unless stated otherwise, components for yeast growth media were purchased from United States Biological. Enzymes and cloning reagents were purchased from New England Biolabs. Genes were purchased from Twist Bioscience.

### 4.2 Cloning

All plasmids used in this study are listed in **Supplementary Table 1**. All primers used for cloning and diagnostic PCR are listed in **Supplementary Table 2**. The sequences of ordered genes are listed in **Supplementary Table 3**. The MoClo Toolkit^38^ and parts from the YTK Toolkit^28^ were used to clone genomic integration cassettes. Genes *matB* and *matC* were ordered with attached BsaI restriction sites and cloned using GoldenGate assembly into Level 1 transcriptional units. pYTK027, pYTK054, and pMYT039 were used to construct transcriptional unit pMATB, and pYTK026, pYTK053, pMYT040 were used to construct transcriptional unit pMATC. pMATB, pMATC, and pMYT075 generated a Level 2 integration cassette using GoldenGate assembly with BsmBI for integration onto the Int. 1 MYT locus on Chromosome I. A KanMX marker was cloned into the Level 2 cassette using Gibson assembly (Primers 1-4), generating pMATBC_1. To generate integration cassettes with *matB*/*matC* promoters of different strengths (pMATBC_2 – pMATBC_5), Gibson assembly using primers 5-22 was used. The *matC* gene was exchanged for *mae1* to generate integration cassettes pMAE1_1 – pMAE1_5 using GoldenGate assembly with primers 23-30. To delete the matC gene from the integration cassette pMATBC_3, GoldenGate assembly with primers 31 and 32 was used to construct pTDH3-MATB.

### 4.3 Strain engineering

All strains used in this study are listed in **Supplementary Table 4**. Integration cassettes pMATBC1 – pMATBC_5, pMAE1_1 – pMAE1_5, and pTDH3-MATB with lanks for homologous integration into the Int. 1 MYT locus on Chromosome I were digested with NotI. The parent strain was transformed with the digested cassettes as described previously^39^. Transformed colonies were selected on plates with added G418 sulfate to select for the KanMX resistance marker (synthetic complete medium: 20 g/L D-glucose, 1.72 g/L yeast nitrogen base w/o AA, carbohydrate and ammonium sulfate, 1 g/L L-glutamic acid, monosodium hydrate salt (Sigma Aldrich), 1.62 g/L dropout-mix synthetic minus leucine w/o yeast nitrogen base, 0.8 mg/ml G418 sulfate). Colony PCR with primers 33-36 was performed to confirm successful integration into the genome, followed by sequencing of the PCR product.

### 4.4 Growth conditions

Yeast cells were grown in liquid synthetic complete growth media containing 20 g/L D-glucose, 1.9 g/L yeast nitrogen base w/o AA, AS, and biotin (Formedium), 5 g/L ammonium sulfate, and 1.62 g/L dropout-mix synthetic minus leucine w/o yeast nitrogen base. In the media with biotin, 2 *μ*g/L biotin was added. In the media with the malonate concentration series, a 500 mM malonic acid disodium salt (Thermo Fisher) solution in water was added to achieve the corresponding concentration. In the media with diethylmalonate concentration series, a 1 M diethylmalonate (Sigma Aldrich) solution in ethanol was added to achieve the corresponding concentration. For passaging experiments aimed at eliminating residual biotin, the yeast cells were incubated in 3 ml cultures in 24-deep-well plates at 30 °C and 300 rpm. For shake-lask experiments, 100 ml media in 250 ml baffled flasks were inoculated with saturated yeast cultures to a starting OD^600^ = 0.01 in triplicates, and the cultivation proceeded at 30 °C and 200 rpm.

### 4.5 Growth curve measurements

Saturated yeast cultures were grown on (diethyl)malonate without biotin and washed 5x with 0.9% NaCl to eliminate residual (diethyl)malonate. The cultures were used to inoculate 200 μl media in a 96-well microtiter plate (Greiner) to achieve a starting OD^600^ of 0.01. Growth was monitored in BMG LabTech CLARIOstar Plus Microplate Reader at 30 °C under agitation (double-orbital shaking, 300 rpm) by measuring the absorbance at 600 nm for 48 h. Each growth curve is an average of 2-4 replicate curves. OD^600^ was calculated from the absorbance based on a calibration curve. The maximum growth rate was calculated using the QurvE software with the parametric growth it^40^.

## 5. Acknowledgements

This project was supported by SNSF 10000664 to T.R.W. and NSF MCB 2438980 to C.C.L. as part of an NSF-SNSF collaborative research program. This project also received funding from the European Union’s Horizon Europe research and innovation program under the Marie Skłodowska-Curie Grant Agreement No.101073546 (Doctoral Network Metal Containing Radical Enzymes – MetRaZymes). Additional funding was provided by the NCCR Molecular Systems Engineering (grant agreement number 200021_178760).

