## Supporting Information for "Biotin-Independent *Saccharomyces cerevisiae* with Enhanced Growth: Engineering an Acetyl-CoA Carboxylase Bypass"

#### Content

1. Supplementary Figures .......... 2
2. Supplementary Tables .......... 7

### 1. Supplementary Figures

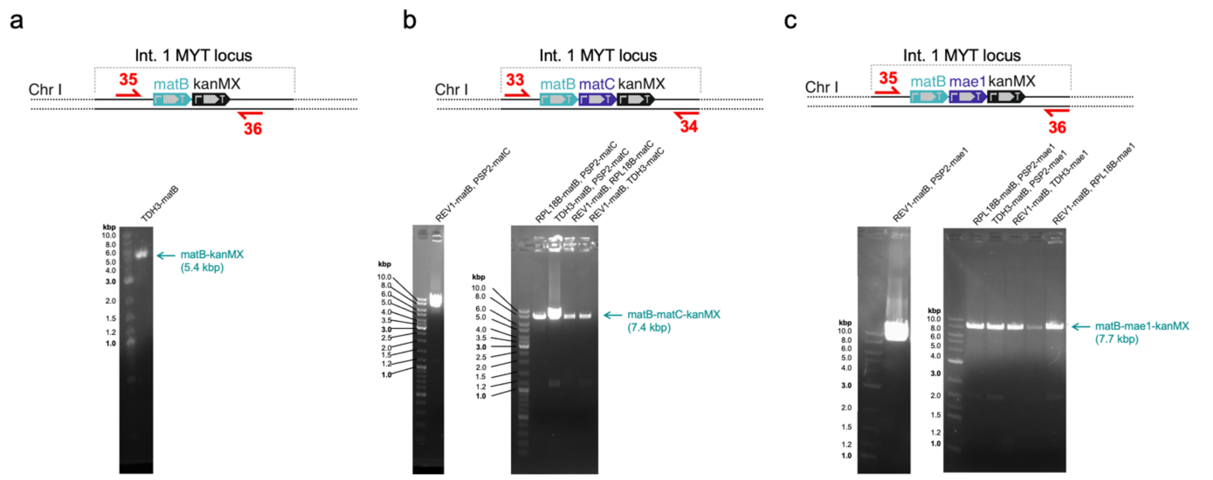


**Supplemetary Fig. 1:** PCR verification of genomic integration of **a)** *matB*, **b)** *matB* and *matC*, and **c)** *matB* and *mae1* genes under the control of weak (REV1, PSP2), medium (RPL18B) or strong (TDH3) promoters into the Int. 1 MYT locus on Chromosome I of *Saccharomyces cerevisiae*. The primers used for colony-PCR are highlighted as red arrows. The products of the PCR reaction were verified by sequencing.

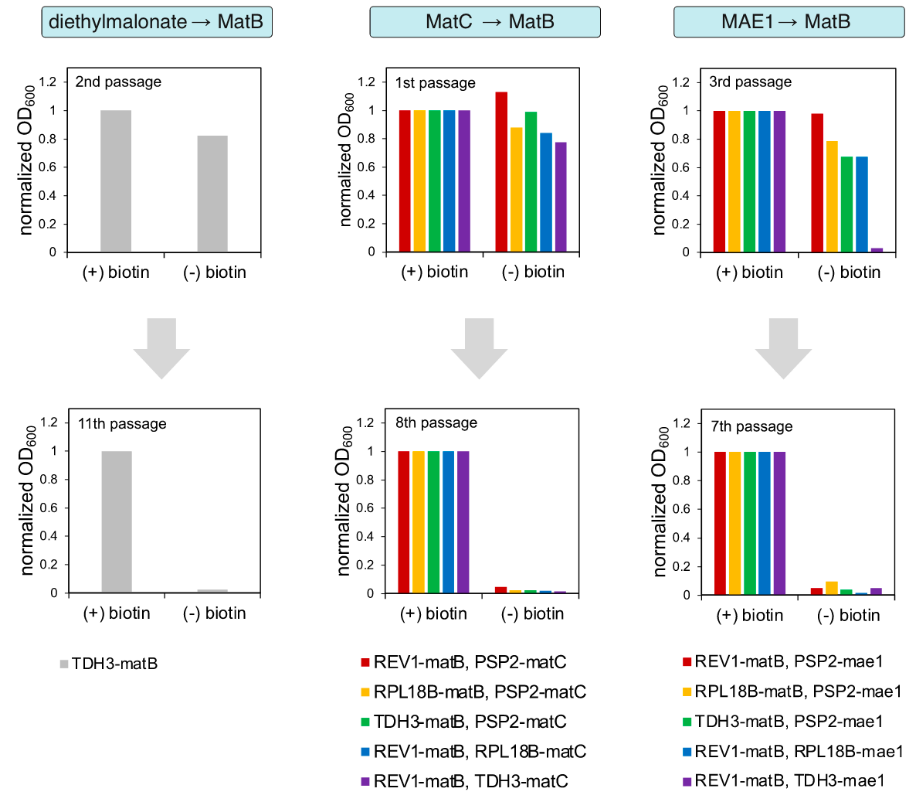


**Supplementary Fig. 2: Passaging of engineered *S. cerevisiae* strains in biotin-free media to eliminate residual biotin.** Each passage corresponds to a 1:1000 dilution of saturated culture into fresh medium. Simultaneously, the strains were passaged in malonate (1-100 mM) or diethyl malonate (1 mM) containing media to generate biotin-free strains for follow-up growth experiments.

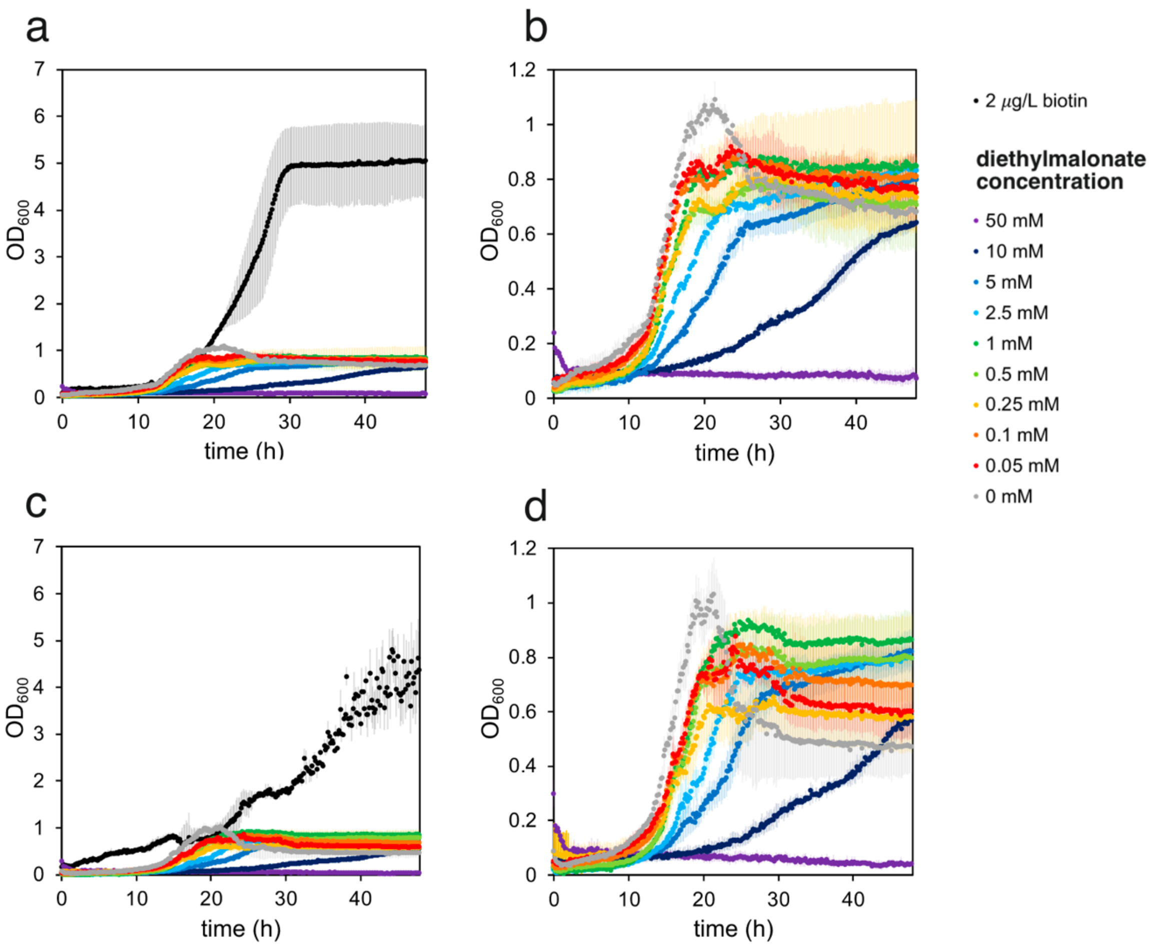


**Supplementary Fig. 3: ACC1 bypass relying on cell-permeable diethyl malonate and MatB.** Growth of the parent strain (**a, b**) and strain with TDH3-*matB* in the Int1 MYT locus (**c, d**) on diethyl malonate concentration series.

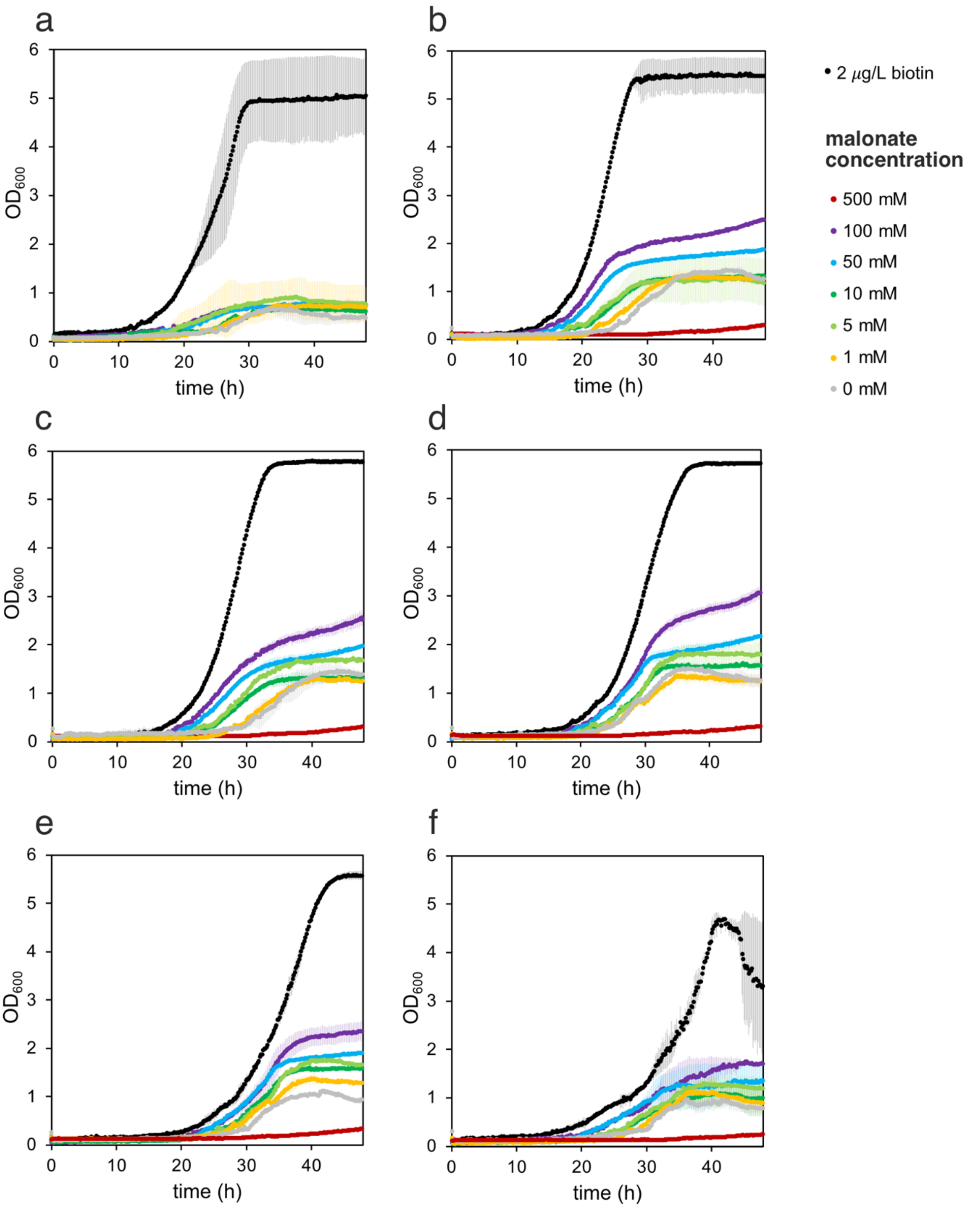


**Supplementary Fig. 4: ACC1 bypass relying on MatC and MatB.** Growth of (**a**) the parent yeast strain, and yeast strains with (**b**) REV1-*matB* and PSP2-*matC*, (**c**) RPL18B-*matB* and PSP2-*matC*, (**d**) TDH3-*matB* and PSP2-*matC*, (**e**) REV1-*matB* and RPL18B-*matC*, and (**f**) REV1-*matB* and TDH3-*matC* in the Int1 MYT locus on malonate concentration series.

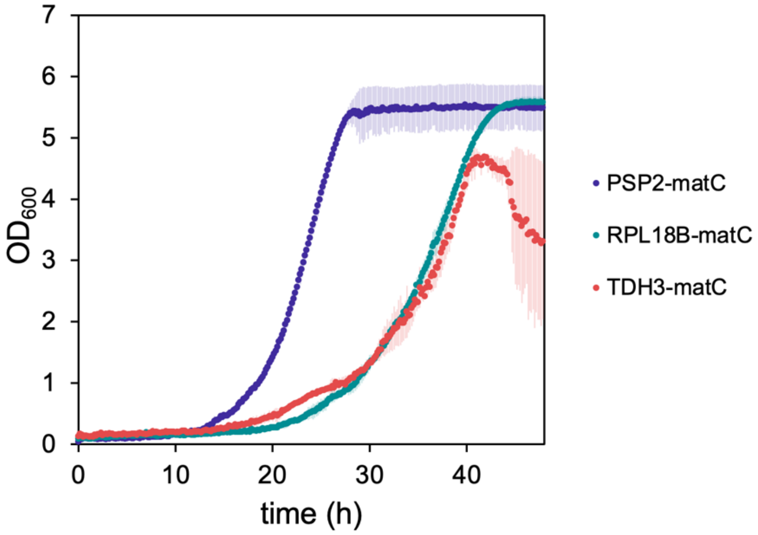


**Supplementary Fig. 5: Toxicity of MatC in *S. cerevisiae*.** Increased expression of MatC in yeast (PSP2 < RPL18B < TDH3) leads to decreased growth and cell death in media containing biotin (2 $\mu$g $L^{-1}$ biotin). All strains contained the *matB* gene under the control of a weak REV1-promoter.

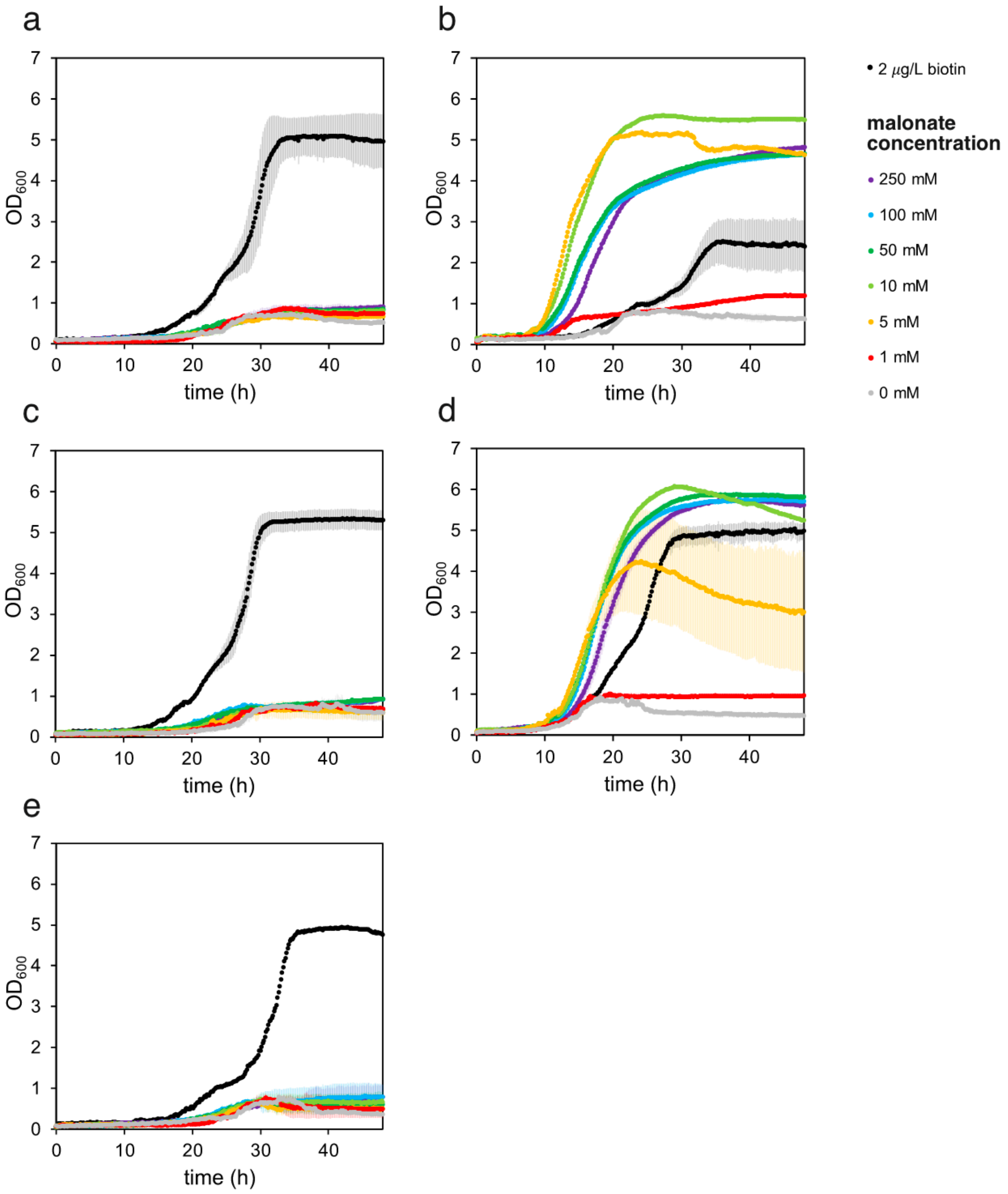


**Supplementary Fig. 6: ACC1 bypass relying on MatC and MAE1.** Growth of yeast strains with (**a**) REV1-*matB* and PSP2-*mae1*, (**b**) RPL18B-*matB* and PSP2-*mae1*, (**c**) TDH3-*matB* and PSP2-*mae1*, (**d**) REV1-*matB* and RPL18B-*mae1*, and (**e**) REV1-*matB* and TDH3-*mae1* in the Int1 MYT locus on malonate concentration series.

### 2. Supplementary Tables

**Supplementary Table 1:** Plasmids used in this study.

| plasmid | description | reference |
|---|---|---|
| pYTK026 | pPSP2 | Lee *et al.*, 2015[1] |
| pYTK027 | pREV1 | Lee *et al.*, 2015[1] |
| pYTK053 | tADH1 | Lee *et al.*, 2015[1] |
| pYTK054 | tPGK1 | Lee *et al.*, 2015[1] |
| pMYT039 | Level 1 assembly cassette | Shaw *et al.*, 2023[2] |
| pMYT040 | Level 1 assembly cassette | Shaw *et al.*, 2023[2] |
| pMYT075 | Int. 1 MYT locus integration vector | Shaw *et al.*, 2023[2] |
| pMATB | pREV1-matB-tPGK1 Level 1 cassette (AmpR) | This study |
| pMATC | pPSP2-matC-tADH1 Level 1 cassette (AmpR) | This study |
| pMATBC_1 | pREV1-matB-tPGK1, pPSP2-matC-tADH1, KanMX Level 2 int. cassette | This study |
| pMATBC_2 | pRPL18B-matB-tPGK1, pPSP2-matC-tADH1, KanMX Level 2 int. cassette | This study |
| pMATBC_3 | pTDH3-matB-tPGK1, pPSP2-matC-tADH1, KanMX Level 2 int. cassette | This study |
| pMATBC_4 | pREV1-matB-tPGK1, pRPL18B-matC-tADH1, KanMX Level 2 int. cassette | This study |
| pMATBC_5 | pREV1-matB-tPGK1, pTDH3-matC-tADH1, KanMX Level 2 int. cassette | This study |
| pMAE1_1 | pREV1-matB-tPGK1, pPSP2-mae1-tADH1, KanMX Level 2 int. cassette | This study |
| pMAE1_2 | pRPL18B-matB-tPGK1, pPSP2-mae1-tADH1, KanMX Level 2 int. cassette | This study |
| pMAE1_3 | pTDH3-matB-tPGK1, pPSP2-mae1-tADH1, KanMX Level 2 int. cassette | This study |
| pMAE1_4 | pREV1-matB-tPGK1, pRPL18B-mae1-tADH1, KanMX Level 2 int. cassette | This study |
| pMAE1_5 | pREV1-matB-tPGK1, pTDH3-mae1-tADH1, KanMX Level 2 int. casette | This study |
| pTDH3-MATB | pTDH3-matB-tPGK1, KanMX Level 2 int. cassette | This study |

**Supplementary Table 2:** Primers used in this study.

| number | sequence (5'-3') |
|---|---|
| 1 | gtcgctatactggcagctaactagtgtacctgcagac |
| 2 | gggcctccatgtctcagcgaaatggggagcg |
| 3 | ccatttcgctgagacatggaggcccagaatacc |
| 4 | actagttagctgccagtatagcgaccagcattcaca |
| 5 | agatctatgtctaaccacctatttgatgcc |
| 6 | cagcattcacatacgattgacgcatga |
| 7 | atatcatgcgtcaatcgtatgtgaatgctggtcgctatactg |
| 8 | attggacatcctcttcgtttcagagtctctcgtttcagagtctctagattgggaattcag |
| 9 | agagactctgaaacgaagaggatgtccaatattttttttaaggaataagg |
| 10 | gttagacatagatcttttgtttttgttttcttctaattgatttttctttctatttcctt |
| 11 | acataaacaaacaaaagatctatgtctaaccacctatttgatgcc |
| 12 | gataaactcgaactgcgtttcagagtctctagattgggaatt |
| 13 | agagactctgaaacgcagttcgagtttatcattatcaatactgc |
| 14 | gttagacatagatcttttgtttgtttatgtgtgtttattcgaaactaagttct |
| 15 | aaaacaaaaaacaaaagatctatgggaattgagctacttagtattgg |
| 16 | attggacatcctcttcgttttggcagcacatagaaatatcgaa |
| 17 | tgtgctgccaaaacgaagaggatgtccaatattttttttaaggaataagg |
| 18 | aattcccatagatcttttgtttttgttttcttctaattgatttttctttctatttcctt |
| 19 | acataaacaaacaaaagatctatgggaattgagctacttagtattgg |
| 20 | gataaactcgaactgcgttttggcagcacatagaaatatcgaa |
| 21 | tgtgctgccaaaacgcagttcgagtttatcattatcaatactgc |
| 22 | aattcccatagatcttttgtttgtttatgtgtgtttattcgaaactaagttct |
| 23 | tatggtctcatgatgggtgaactcaaggaaatcttgaaac |
| 24 | tatggtctcagatttaaacgctttcatgttcactactaggagg |
| 25 | tatggtctcaatcatagatctttcttcccgttccagcactct |
| 26 | tatggtctcaaatcctaactcgaggcgaatttcttatg |
| 27 | tatggtctcaatcacatagatctttcttcccgttccagc |
| 28 | tatggtctcaaatcatcctaactcgaggcgaatttcttatga |
| 29 | tatggtctcaatcacatagatcttttgtttttgttttcttctaattgatttttctttc |
| 30 | tatggtctcaatcacatagatcttttgtttgtttatgtgtgtttattcgaaac |
| 31 | tatggtctcaaacgcagttcgagtttatcattatcaatact |
| 32 | tatggtctcacgtttcagagtctctagattgggaat |
| 33 | ccgaagttcatatacgaatgc |
| 34 | gcattgcaagatattgaataactg |

| | |
|---|---|
| 35 | aggaagtctcgctaagtagg |
| 36 | tgggcatacctgcaatatga |

**Supplementary Table 3: Synthetic genes used in this study**. Overhangs on *matB* and *matC* genes for GoldenGate assembly are highlighted in bold.

| name | sequence (5'-3') |
|---|---|
| *matB* | **gcatcgtctcatcggtctcat**atgtctaaccacctatttgatgccatgcgtgcggcagcaccgggaaatgcaccattcattcgtatcgac<br>aatactagaacgtggacctacgacgacgcgttcgcactaagtggtaggattgcgtctgctatggatgccttaggaataagacccggtgat<br>agggtcgccgttcaagtcgaaaagagtgcggaagctctgatactgtatctggcctgtcttaggtccggggccgtttaccttccgctaaata<br>cagcttataccctagctgagttagattactttataggtgatgctgagcctcgtctagtggtagtagcctccagcgcgagggctggtgtcga<br>gacaatcgccaagccgcgtggagctatagtagaaactttagatgcggcaggctcaggctcccttttagacttagccagggacgagcccg<br>ccgactttgtggatgcgagtaggagtgctgacgatctggcggctattctatatacttctggcacgaccggacgtagcaaaggagcaatgc<br>taacgcacggcaacttgttgtcaaacgccttaactctgagagacttttggagagtaactgctggtgacaggctgatccatgcgctgcctat<br>atttcacacgcacgggttgtttgttgctactaatgtcacacttctagctggcgccagtatgttccttttaagcaaattcgatcccgaagagat<br>actaagtctgatgccacaagcaacgatgttaatgggagttccaaccttttacgtccgtctattgcaatcaccgaggcttgacaaacaagcg<br>gtggctaatatcaggctttttatctccgggtctgcgcctcttctagctgaaacgcacacagagtttcaagcaagaacaggacatgcaatat<br>tggagcgttatggcatgacagaaacaaacatgaacacgagtaatccttacgaaggtaaaaggattgcgggcactgtaggcttcccctta<br>cctgatgtcactgtgagagttactgatcctgcgacgggacttgcactgcctcccgaacagaccgggatgatcgagatcaaaggtcctaac<br>gtcttcaagggttactggaggatgccggaaaaaaccgctgcggaattcacggctgatggtttttttataagtggcgatttaggcaaaatcg<br>atcgtgacggatacgtccacattgttggcaggggggaaggatcttgtgatctctggaggttataatatctatcccaaagaagttgagggtga<br>aatcgatcaaattgaaggggtggtggaaagtgcggttataggggtcccccacccggacttcggagaaggagtaactgcggtagtagta<br>agaaagcccggggcagccctagatgagaaagcgattgtttcagcattacaggacagattagccaggtataagcagccaaagcgtataa<br>tctttgctgaagatttgccaagaaatactatggggaaagtacaaaagaacattcttagacaacaatatgcagatttgtacacacgtacgta<br>**atcctgagacctgagacggcat** |
| *matC* | **gcatcgtctcatcggtctcat**atgggaattgagctacttagtattggtcttttgatcgctatgtttatcatagcaactatccaaccaattaat<br>atgggagctctggcatttgctggggcgttcgtgctggggagcatgattattggaatgaagactaatgaaatctttgcagggttcccaagtg<br>acctatttctgaccttggtagcggtgacgtacttatttgccattgcccaaatcaacggtaccatcgattggctggttgaatgtgcggtgaga<br>ctagtaagaggcagaatagggttaataccttgggtaatgtttctggttgcagccataattactgggtttggtgcgctaggcccagccgccg<br>tagcgatccttgcgcccgttgcgctgagctttgccgtccagtataggatccacccagttatgatgggacttatggtaatccatggggcgca<br>ggcgggcgggttcagtcccatatctatttacggagggataactaatcagatagtggccaaggctgggctaccctttgcaccaacatcact<br>gtttttgtcttctttttctttaacctagcgatagcagtgttggtgttcttcgtattcggtggagctagagttatgaagcatgatcccgcttccct<br>ggggccactgccagagctacaccctgaaggagtaagcgctagtatcagaggccatggcggaaccccagccaagccgatcagggaaca<br>cgcgtatggtacagccgccgacacagccacgacattgagactgaacaacgaaagaatcactacgcttataggcctgacagcccttggg<br>attggcgctttagtgtttaaatttaacgtaggtttagtggctatgacagtcgcagttgtactagccctgctatcacccaaaacgcagaaagc<br>tgcgattgataaggtttcttggtccaccgtattgctgatagcggggattataacttatgtaggtgtcatggaaaaggccggcacggttgact<br>acgtagctaatggtatttcctccttgggaatgccgctattggttgcgctacttctatgcttcacgggagcgattgtctctgcgtttgcctccag<br>tacagcgttgctaggcgctataataccccctggcggtgccgtttctgctgcaaggccatatctccgcaattggcgttgtcgctgccattgcca<br>taagtaccacgattgtggacacgtcaccttttagcactaacggggccttggtggtagcaaacgcacctgacgattctcgtgaacaagtctt<br>gcgtcagctgctaatttatagcgcgttgatcgcaatcatcgggcccattgttgcgtggttagtttttgtggtgccaggattggtttaa**atcct<br>gagacctgagacggcat** |
| *mae1* | Atgggtgaactcaaggaaatcttgaaacagaggtatcatgagttgcttgactggaatgtcaaagcccctcatgtccctctcagtcaacga<br>ctgaagcattttacatggtcttggtttgcatgtactatggcaactggtggtgttggtttgattattggttctttcccctttcgattttatggtctta<br>atacaattggcaaaattgtttatattcttcaaatctttttgttttctctctttggatcatgcatgctttttcgctttattaaatatccttcaactatc<br>aaggattcctggaaccatcatttggaaaagcttttcattgctacttgtcttctttcaatatccacgttcatcgacatgcttgccatatacgcct<br>atcctgataccggcgagtggatggtgtgggtcattcgaatcctttattacatttacgttgcagtatcctttatatactgcgtaatggctttttt<br>acaattttcaacaaccatgtatataccattgaaaccgcatctcctgcttggattcttcctattttccctcctatgatttgtggtgtcattgctgg<br>cgccgtcaattctacacaacccgctcatcaattaaaaaatatggttatctttggtatcctctttcaaggacttggtttttgggtttatcttttac<br>tgtttgccgtcaatgtcttacggtttttactgtaggcctggcaaaaccccaagatcgacctggtatgtttatgtttgtcggtccaccagcttt<br>ctcaggtttggccttaattaatattgcgcgtggtgctatgggcagtcgcccttatatttttgttggcgccaactcatccgagtatcttggttttg<br>tttctacctttatggctatttttatttggggtcttgctgcttggtgttactgtctcgccatggttagctttttagcgggctttttcactcgagcccc<br>tctcaagtttgcttgtggatggtttgcattcattttccccaacgtgggttttgttaattgtaccattgagataggtaaaatgatagattccaaa<br>gctttccaaatgtttggacatatcattggggtcattctttgtattcagtggatcctcctaatgtatttaatggtccgtgcgtttctcgtcaatga<br>tctttgctatcctggcaaagacgaagatgcccatcctccaccaaaaccaaatacaggtgtccttaaccctaccttcccacctgaaaaagca<br>cctgcatctttggaaaaagtcgatacacatgtcacatctactggtggtgaatcggatcctcctagtagtgaacatgaaagcgtttaa |

**Supplementary Table 4:** Yeast strains used in this study.

| name | description | source |
|---|---|---|
| parent strain | F102-2 Leu2△0 ura3△0 His4+ his3△1 flo1△0 met15△0 trp5△0 can1::pPSP2-TPDNAP1-tADH1_NatMX pGKL1::Leu2 | UC Irvine, CA |
| TDH3-*matB* | F102-2 Leu2△0 ura3△0 His4+ his3△1 flo1△0 met15△0 trp5△0 can1::pPSP2-TPDNAP1-tADH1_NatMX pGKL1::Leu2 Int1:: pTDH3-matB-tPGK1_KanMX | This study |
| REV1-*matB*, PSP2-*matC* | F102-2 Leu2△0 ura3△0 His4+ his3△1 flo1△0 met15△0 trp5△0 can1::pPSP2-TPDNAP1-tADH1_NatMX pGKL1::Leu2 Int1:: pREV1-matB-tPGK1_pPSP2-matC-tADH1_KanMX | This study |
| RPL18B-*matB*, PSP2-*matC* | F102-2 Leu2△0 ura3△0 His4+ his3△1 flo1△0 met15△0 trp5△0 can1::pPSP2-TPDNAP1-tADH1_NatMX pGKL1::Leu2 Int1:: pRPL18B-matB-tPGK1_pPSP2-matC-tADH1_KanMX | This study |
| TDH3-*matB*, PSP2-*matC* | F102-2 Leu2△0 ura3△0 His4+ his3△1 flo1△0 met15△0 trp5△0 can1::pPSP2-TPDNAP1-tADH1_NatMX pGKL1::Leu2 Int1:: pTDH3-matB-tPGK1_pPSP2-matC-tADH1_KanMX | This study |
| REV1-*matB*, RPL18B-*matC* | F102-2 Leu2△0 ura3△0 His4+ his3△1 flo1△0 met15△0 trp5△0 can1::pPSP2-TPDNAP1-tADH1_NatMX pGKL1::Leu2 Int1:: pTDH3-matB-tPGK1_pRPL18B-matC-tADH1_KanMX | This study |
| REV1-*matB*, TDH3-*matC* | F102-2 Leu2△0 ura3△0 His4+ his3△1 flo1△0 met15△0 trp5△0 can1::pPSP2-TPDNAP1-tADH1_NatMX pGKL1::Leu2 Int1:: pTDH3-matB-tPGK1_pTDH3-matC-tADH1_KanMX | This study |
| REV1-*matB*, PSP2-*mae1* | F102-2 Leu2△0 ura3△0 His4+ his3△1 flo1△0 met15△0 trp5△0 can1::pPSP2-TPDNAP1-tADH1_NatMX pGKL1::Leu2 Int1:: pREV1-matB-tPGK1_pPSP2-mae1-tADH1_KanMX | This study |
| RPL18B-*matB*, PSP2-*mae1* | F102-2 Leu2△0 ura3△0 His4+ his3△1 flo1△0 met15△0 trp5△0 can1::pPSP2-TPDNAP1-tADH1_NatMX pGKL1::Leu2 Int1:: pRPL18B-matB-tPGK1_pPSP2-mae1-tADH1_KanMX | This study |
| TDH3-*matB*, PSP2-*mae1* | F102-2 Leu2△0 ura3△0 His4+ his3△1 flo1△0 met15△0 trp5△0 can1::pPSP2-TPDNAP1-tADH1_NatMX pGKL1::Leu2 Int1:: pTDH3-matB-tPGK1_pPSP2-mae1-tADH1_KanMX | This study |
| REV1-*matB*, RPL18B-*mae1* | F102-2 Leu2△0 ura3△0 His4+ his3△1 flo1△0 met15△0 trp5△0 can1::pPSP2-TPDNAP1-tADH1_NatMX pGKL1::Leu2 Int1:: pTDH3-matB-tPGK1_pRPL18B-mae1-tADH1_KanMX | This study |
| REV1-*matB*, TDH3-*mae1* | F102-2 Leu2△0 ura3△0 His4+ his3△1 flo1△0 met15△0 trp5△0 can1::pPSP2-TPDNAP1-tADH1_NatMX pGKL1::Leu2 Int1:: pTDH3-matB-tPGK1_pTDH3-mae1-tADH1_KanMX | This study |
